# Experimental Investigation of Pulse Sterilization of Pathogens

**DOI:** 10.1101/2021.11.25.470057

**Authors:** Volodymyr Chumakov, Mykhailo Ostryzhnyi, Oksana Kharchenko, Nataliya Rybalchenko, Vasiliy Muraveinyk, Aleksandr Tarasevich

## Abstract

The results of experimental investigations of the effect of high-intensity pulsed UV radiation on reference strains of opportunistic pathogens S, aureus and E. coli are presented. The modified pulse UV sterilizer MПК-300-3 based on an end-face plasma accelerator was used as a radiation source, which provides a power pulsed discharge in an open atmosphere. The high efficiency of inactivation of the pathogens was provided within a short period of time. The possibility of providing urgent 100% sterilization of a pathogens has been shown. The prospects for the application of pulse sterilization technology to combat pathogens are considered.

## INTRODUCTION

Research on the mechanism and effectiveness of pulsed UV sterilization is ongoing. As a matter of fact, they did not stop, having started almost 30 years ago [1-5]. Successful tests of the technology of pulse sterilization of viral infection opened the prospect for further research on the effect of UV radiation on microorganisms [6]. The results of all the studies carried out show that the high efficiency of pulse sterilization is provided, first of all, by the high power of UV radiation, which is generated in the pulsed mode. At the same time, generators of optical radiation based on magnetoplasmatic compressor (MPC) are the most powerful emitting devices. The purpose of these studies was to show the high efficiency of incoherent pulsed UV-radiation generated during the discharge of an end-face coaxial MPC to pathogens and to experimentally confirm the possibility of monopulse sterilization.

## 1. MATERIALS AND METHODS

Studies were performed on reference strains of opportunistic pathogens *Staphylococcus aureus* UCM B-904 (ATCC 25923), *Escherichia coli* UCM B-906 (ATCC 25922), obtained from the Ukrainian collection of microorganisms (UCM, Institute of Microbiology and Virology of NAS of Ukraine) [7].

To determine the sensitivity of microorganisms to UV-radiation, a test culture of bacteria was incubated for 24 hours at 37^0^ C on solid medium LB (Luria-Bertoni). Daily agar cultures were washed with sterile saline, diluted with a suspension according to the McFarland 1.0 turbidity standard, followed by dilution to 1×10^6^ CFU (colony-forming units)/ml, 1×10^4^ CFU/ml, 1×10^3^ CFU/ml [8]. Volume 0.1 ml bacterial suspension of the appropriate concentration was inoculated by the lawn on Mueller-Hinton agar plates. Source of UV-radiation was placed so that the device was directed downward and placed at a height of *h =* 30 cm from the surface of the laboratory table (Fig. 1). On the table under the ultraviolet sterilizer in the operation area the opened Petri cups inoculated with a certain concentration of bacterial suspension were placed, and a single dose of radiation (1 pulse) was exposed. Next, the irradiated cups were closed and set aside, and in their place the next series of inoculated cups were placed and two exposures of radiation (2 pulses) were given. The same principle was used to irradiate 5 and 10 pulses. Petri cups not irradiated with an UV-radiation served as a control. The experiment used three replicates of inoculated cups with different concentrations of bacterial suspension. Irradiated and control cups were incubated in a thermostat at 37° C for 24 hours.

**Fig. 1.**
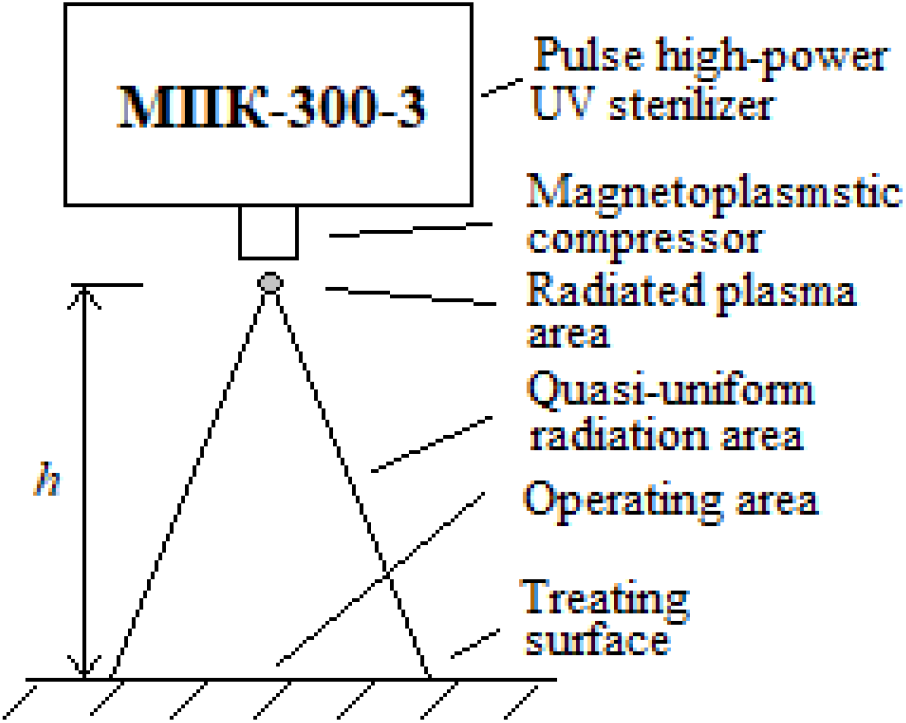
Experimental investigation scheme

The dishes inoculated with cultures in different concentrations were divided into 3 groups, each of which was irradiated with a certain dose of UV radiation. The dose load on a group of cups was set by the number of pulsed exposures

When calculating the results, the number of viable cells was determined, the number of colonies grown on the plates was counted, the cell titer and the survival of irradiated cells as a percentage of control were determined.

A modified MPK-300-3 sterilizer was used as a source of pulsed UV radiation [6]. The circuitry implementation of the main units of the device, the control circuit and the appearance have undergoned modernization. The switching elements of the charging-discharge circuit of the electric energy storage have been completely renewed and strengthened. Fig. 2 represents an updated equipment. Currently, the device provides operation both in the mode of generating single pulses and in the mode of forming a pulse train of a given duration. The use of modern element base has made it possible to increase the stability of the parameters of radiation pulses.

**Fig. 2.**
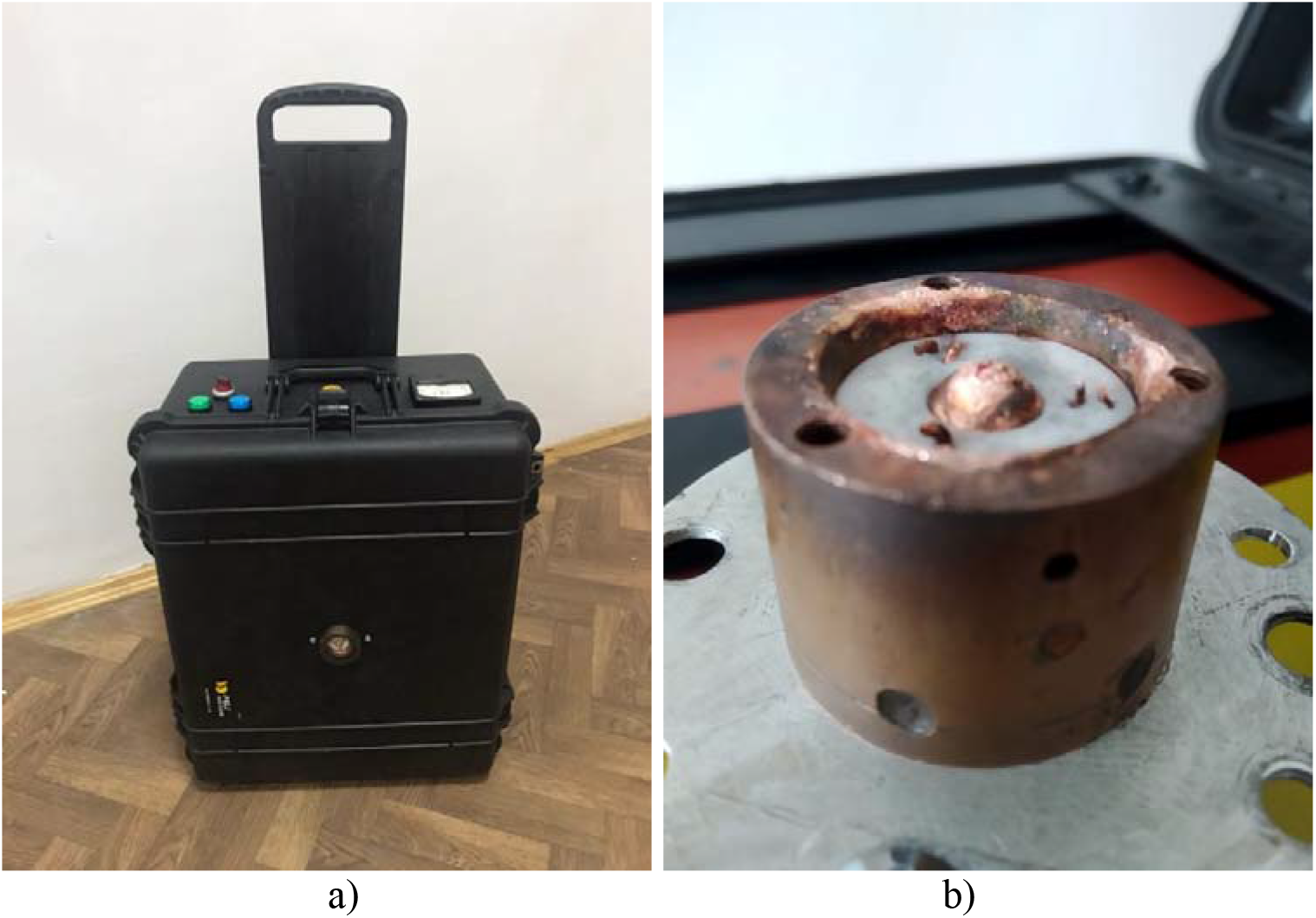
Updated model of the high-power pulse UV sterilizer MПК-300-3 (a), radiator unit (MPC) (b)

## 2. RESULTS AND DISCUSSION

The results of the study of the effectiveness of the pulsed UV-radiation against reference strains of opportunistic pathogens *<u>S. aureus</u>* and *E*.*coli* are presented in Tab. 1, 2 and Fig. 3 – 8.

**Table 1.** The effectiveness of pulsed UV-radiation against *S. Aureus*

| Test-culture | Initial bacterial suspension concentration, control, CFU/ml | Number of pulse exposition | Resulted number of growth colonies | Efficiency % |
| --- | --- | --- | --- | --- |
| *S. aureus* | $10^6$ | 1 | 272 | 99,97 |
|  |  | 2 | 124 | 99,98 |
|  |  | 5 | 16 | 99,99 |
|  |  | 10 | 0 | 100 |
| | $10^4$ | 1 | 35 | 99,65 |
|  |  | 2 | 20 | 99,80 |
|  |  | 5 | 2 | 99,98 |
|  |  | 10 | 0 | 100 |
| | $10^3$ | 1 | 1 | 99,90 |
|  |  | 2 | 1 | 99,90 |
|  |  | 5 | 0 | 100 |
|  |  | 10 | 0 | 100 |

**Table 2.** The effectiveness of pulsed UV-radiation against *E. coli*

| Test-culture | Initial suspension concentration, control, CFU/ml | Number of pulse exposition | Resulted number of growth colonies | Efficiency % |
| --- | --- | --- | --- | --- |
| *E. coli* | 10<sup>6</sup> | 1 | 468 | 99,95 |
|  |  | 2 | 71 | 99,99 |
|  |  | 5 | 13 | 99,99 |
|  |  | 10 | 0 | 100 |
|  | 10<sup>4</sup> | 1 | 68 | 99,32 |
|  |  | 2 | 7 | 99,93 |
|  |  | 5 | 0 | 100 |
|  |  | 10 | 0 | 100 |
|  | 10<sup>3</sup> | 1 | 10 | 99,00 |
|  |  | 2 | 2 | 99,80 |
|  |  | 5 | 0 | 100 |
|  |  | 10 | 0 | 100 |

**Fig. 3.**
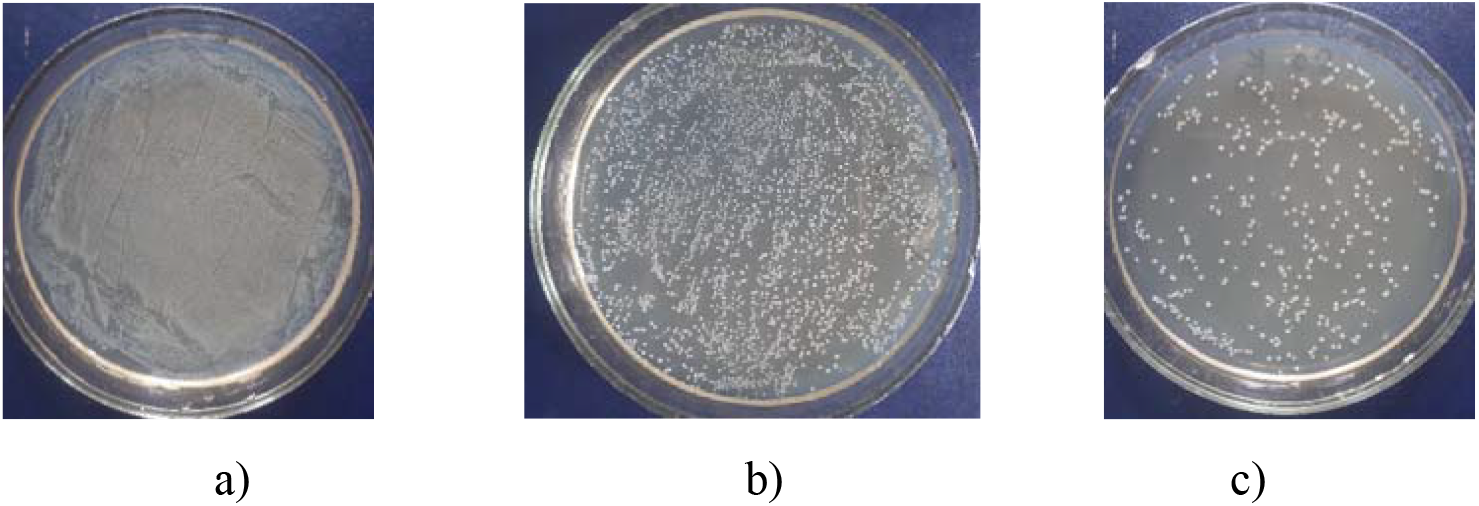
Initial inoculated cups with *S. aureus N*_0_ = 10^6^ CUO/ml (a), *N*_0_ = 10^4^ CUO/ml (b) and *N*_0_ = 10^3^ CUO/ml (c)

**Fig. 4.**
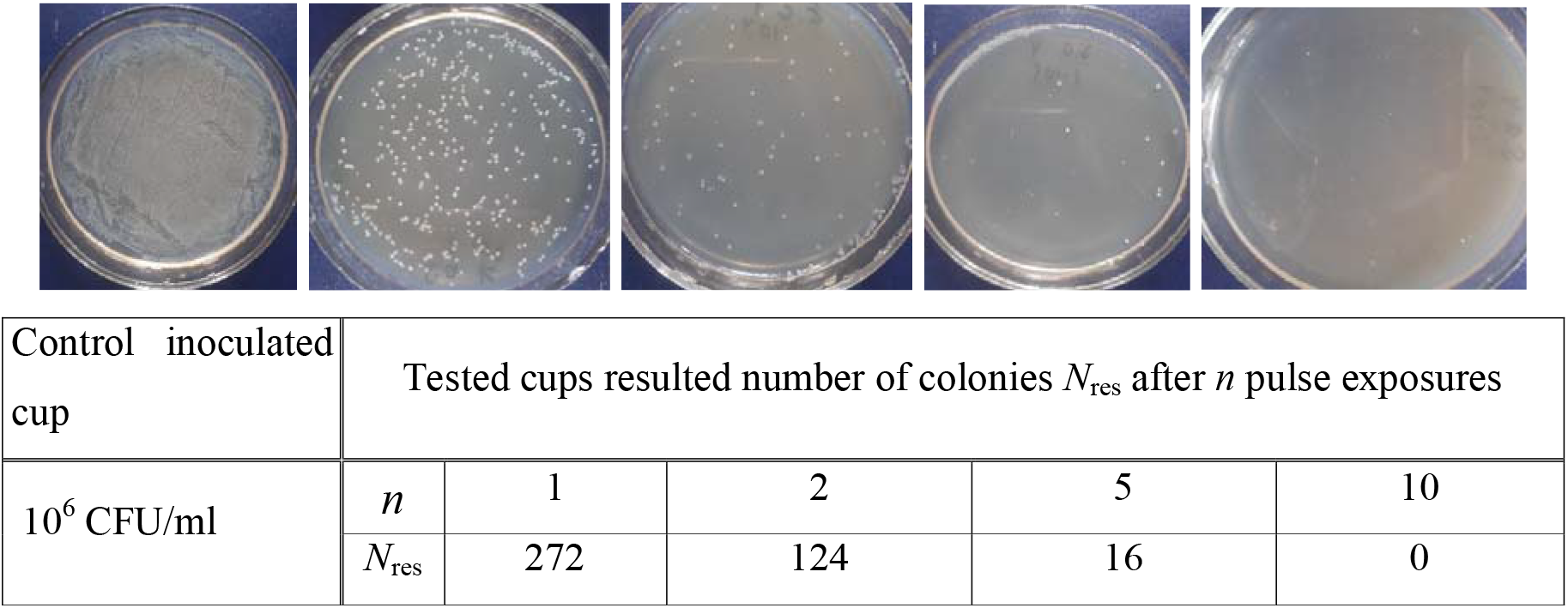
Control and irradiated Petri cups inoculated with *S*.*aureus* and resulted number colonies dependence on pulse exposition

**Fig.5.**
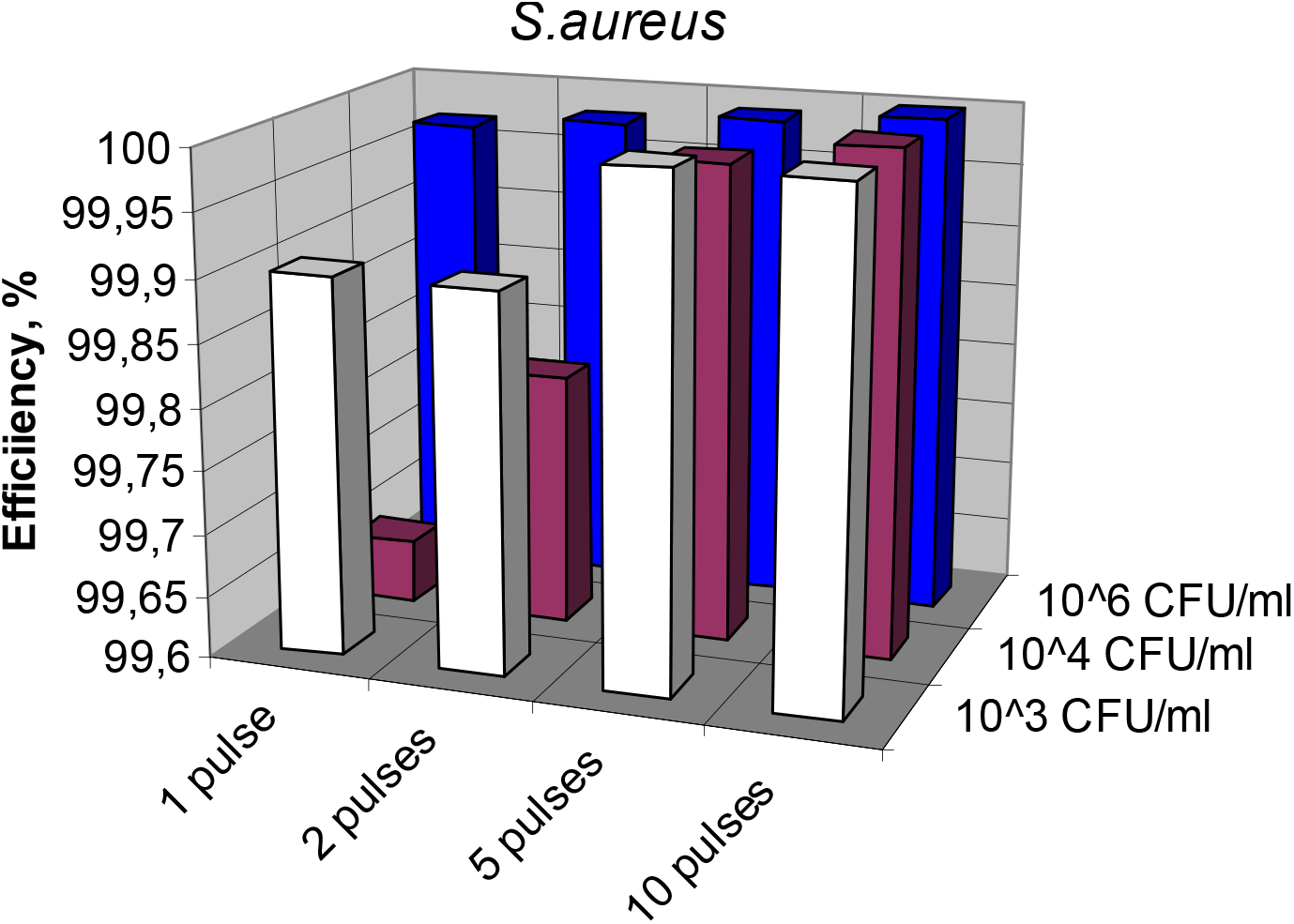
*S. aureus* sterilization efficiency dependence on number of pulse exposition

**Fig. 6.**
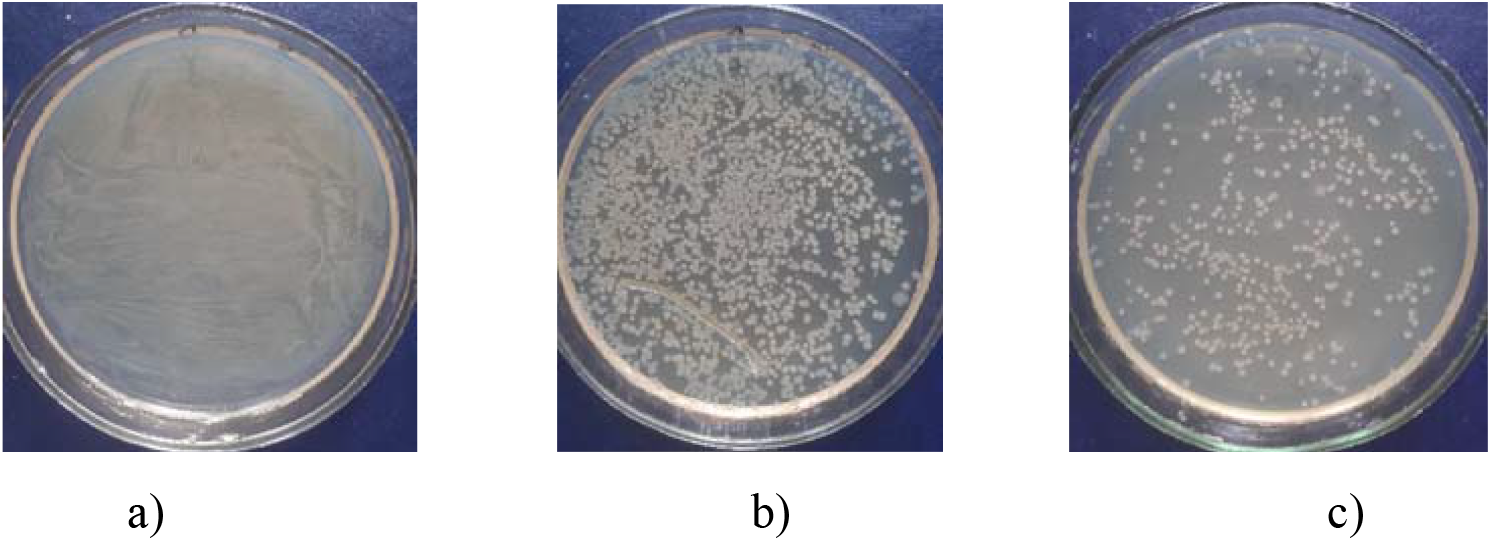
Initial inoculated cups with *E. coli N*_0_ = 10^6^ CUO/ml (a), *N*_0_ = 10^4^ CUO/ml (b) and *N*_0_ = 10^3^ CUO/ml (c)

**Fig. 7.**
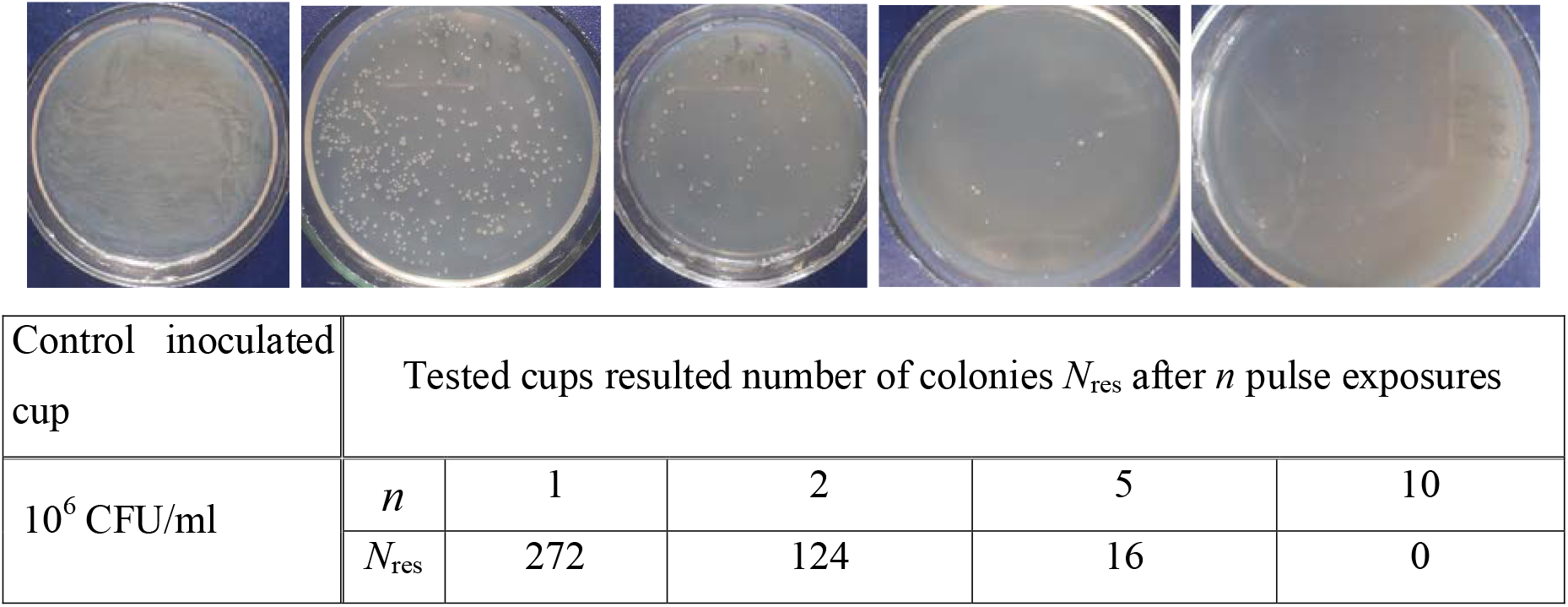
Control and irradiated Petri cups inoculated with *E. coli* and resulted number colonies dependence on pulse exposition

**Fig.8.**
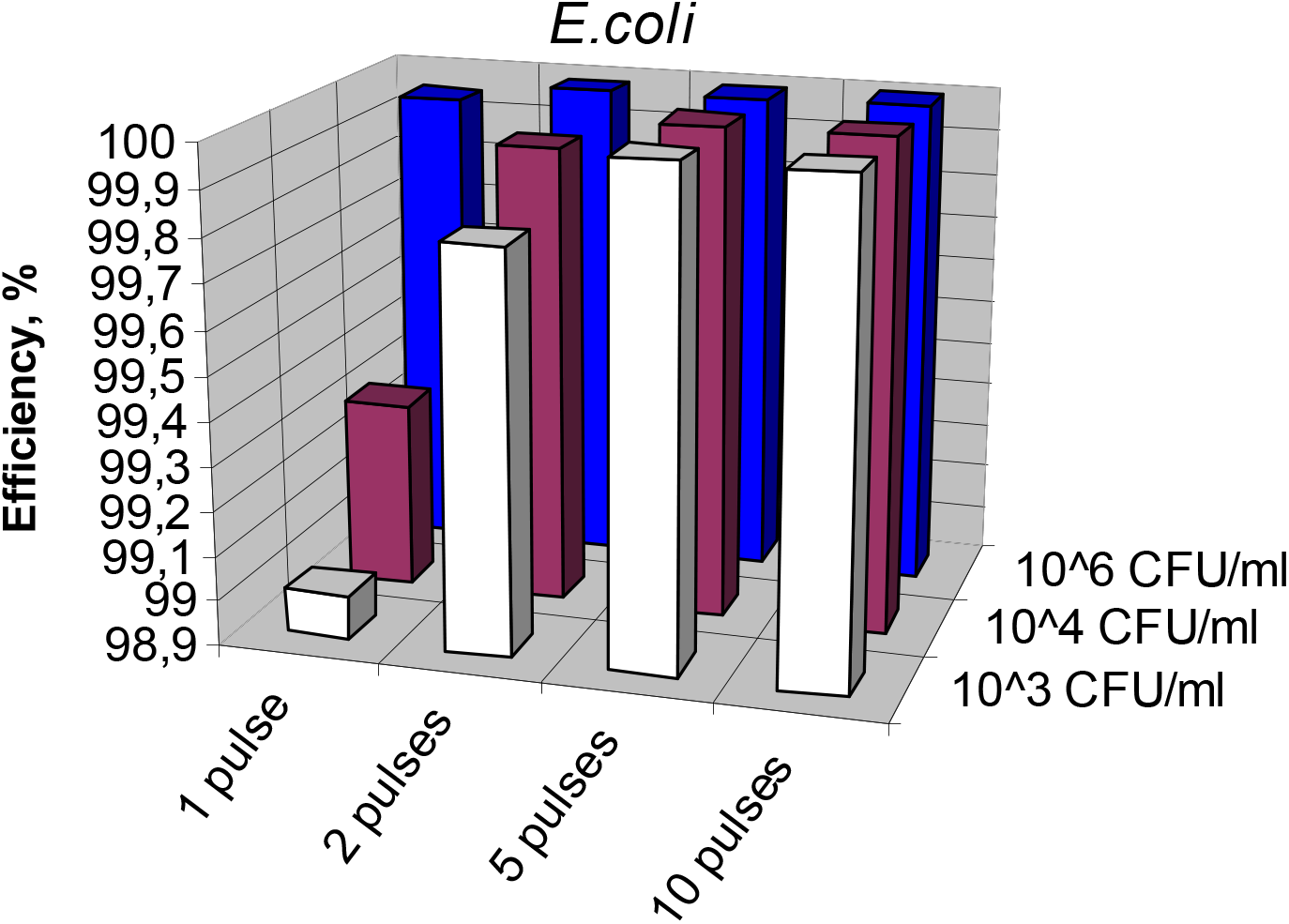
*E*.*coli* sterilization efficiency dependence on number of pulse exposition

As can be seen from the results of the research, the ultraviolet sterilizer MПK-300-3 showed high bactericidal activity against both test cultures of microorganisms. Already under the action of 1 pulse of the sterilizer MПK-300-3 registered inhibition of growth of *S. aureus* by 99.97% at a concentration of bacterial suspension of 1×10^6^ CFU/ml, 99.65% –1×10^4^ CFU/ml, 99.90% – 1×10^3^ CFU/ml. Under the influence of 2 and 5 pulses there is also a high efficiency of the device reaching 99.80 – 99.99%, respectively. Under the influence of 10 pulses of the sterilizer, 100% death of *S. aureus* cells is detected in all studied concentrations of bacterial suspension. We also note that at an initial concentration of pathogens equal to 1×10 CFU/ml, the microbial residue after a single exposure was only 1 CFU, which is almost equivalent to the effect of monopulse sterilization.

Inhibition of E. coli cell growth is also registered under the influence of 1 pulse of sterilizer MПK-300-3 by 99.95% at a concentration of bacterial suspension of 1×10^6^ CFU/ml, 99.32% – 1×10^4^ CFU/ml, 99.00 % – 1×10^3^ CFU/ml. Under the influence of 2 pulses, the efficiency of the device reaches 99.80 – 99.99%. Under the action of 5 pulses of the sterilizer, 100% death of E. coli cells is detected in all studied concentrations of bacterial suspension (Tab. 2). Thus, *E. coli* bacteria are less resistant to pulsed UV radiation.

Therefore, based on the obtained data, it was found that the portable pulsed ultraviolet sterilizer MПK-300-3 has a high bactericidal effect on the studied opportunistic microorganisms *S. aureus* and *E. coli* with an efficiency of 100%, and can be recommended for use to disinfect surfaces, products, premises of various function.

## CONCLUSIONS

1. A pulsed source of high-power UV radiation has been developed and created for analyzing the bactericidal effect and determining the effectiveness of pulse sterilization.
2. The high bactericidal efficiency of high-power pulsed UV radiation when exposed to pathogens has been experimentally shown.
3. Exposure to pulsed UV radiation results in 100% complete sterilization as a result of irradiation with a finite number of pulsed exposures. Thus, the complete destruction of opportunistic pathogenic microorganisms S. aureus is achieved as a result of exposure to 10 pulsed exposures. The complete destruction of S. coli bacteria is achieved as a result of exposure to 5 pulses. A decrease in the initial concentration leads to a decrease in the pulsed radiation dose required for complete sterilization.
4. The results obtained determine the fundamental possibility of monopulse sterilization.

